# Fission yeast obeys a linear size law under nutrient titration

**DOI:** 10.1101/2023.04.12.536544

**Authors:** François Bertaux, Istvan T. Kleijn, Samuel Marguerat, Vahid Shahrezaei

## Abstract

Steady-state cell size and geometry depend on growth conditions. Here, we use an experimental setup based on continuous culture and single-cell imaging to study how cell volume, length, width and surface-to-volume ratio vary across a range of growth conditions including nitrogen and carbon titration, the choice of nitrogen source, and translation inhibition. Overall, we find cell geometry is not fully determined by growth rate and depends on the specific mode of growth rate modulation. However, under nitrogen and carbon titrations, we observe that the cell volume and the growth rate follow the same linear scaling.

Figure
**A**. Graphical outline of the growth and imaging assays. **B**. Illustration of the procedure used to extract cell size and geometry data. **C**. Average surface-to-volume (S/V) ratio plotted as a function of average cell width across all steady-state cultures. The dark grey circle indicates the base growth medium, EMM2. Cultures limited in their growth by the concentration of ammonium, glucose, and cycloheximide in the medium are indicated with cyan lozenges, green triangles, and orange squares, respectively. Cultures limited by the choice of nitrogen source are indicated with light grey circles; the amino-acid nitrogen source used is labelled with its three-letter abbreviation, using Amm for the equivalent culture grown with ammonium chloride as its sole nitrogen source. **D**. Average cell volume plotted against the growth rate across all cultures, showing collapse for ammonium- and glucose-limited cultures and differing behaviour for nitrogen-source- and translation-limited cultures. Plotted in dark grey is a linear fit to the ammonium- and glucose-limited data, including the base medium, with 95% confidence interval; a similar fit (without CI) to the cycloheximide-limited cultures is indicated by a dashed orange line. **E**. Average surface-to-volume (S/V) ratio against growth rate across all cultures, with dashed lines for ammonium-, glucose-, and translation-limited cultures representing linear fits to the respective data, each including the base medium. Under ammonium limitation, the surface-to-volume ratio increases markedly as the growth rate decreases. A moderate increase is observed for glucose limitation and a moderate decrease is observed for protein synthesis inhibition with cycloheximide. There is no consistent trend with the growth rate when the quality of the nitrogen source is varied. **F**. Average surface area, **G**. cell length, **H**. cell width against growth rate, showing that ammonium-limited cells are thinner than glucose-limited cells at equivalent growth rates, consistent with their different surface-to-volume ratios and the relation between surface-to-volume ratio and cell width (see C).

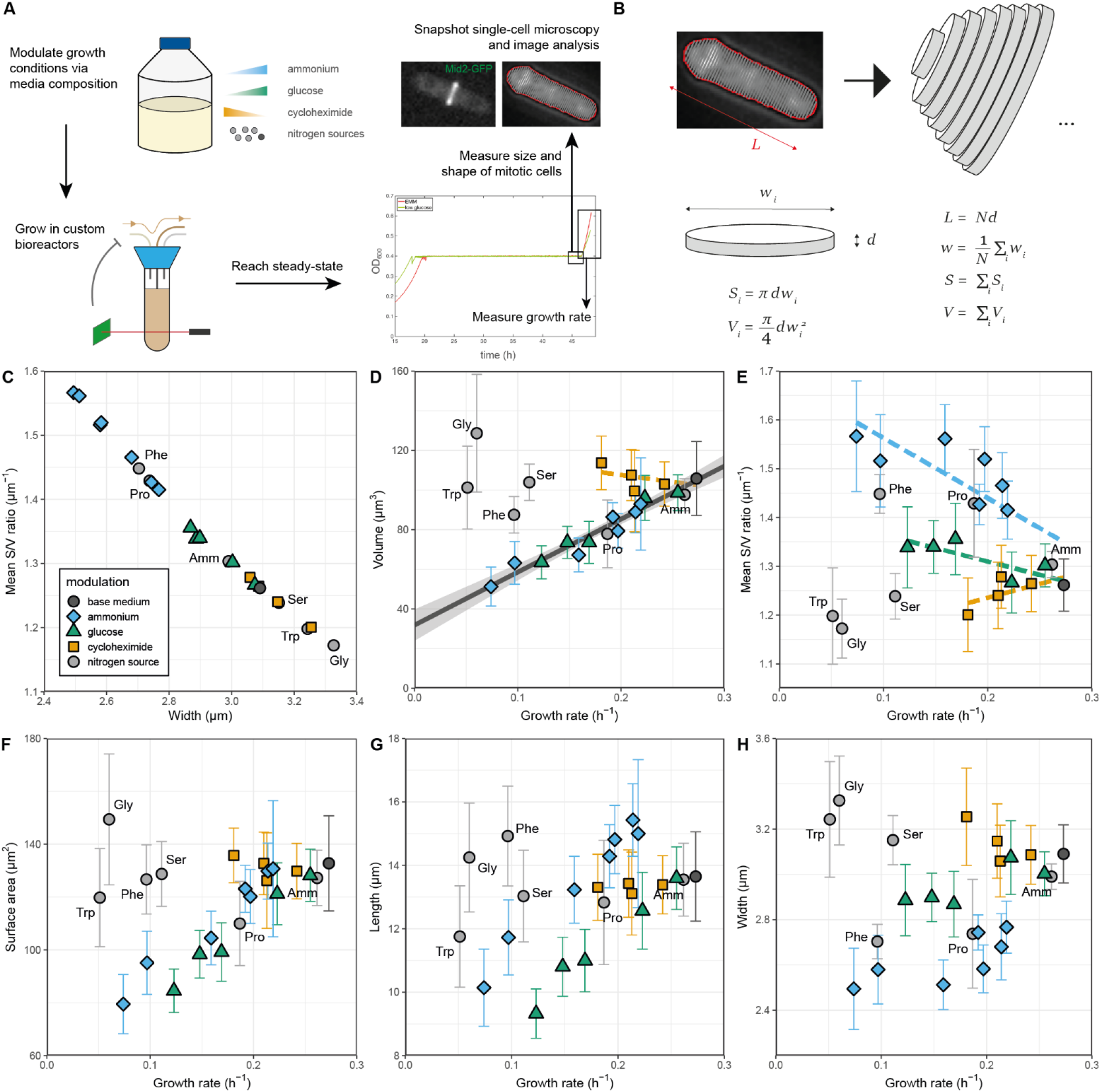

## Description

It has been long known that bacterial cell size increases with cellular growth rate (Schaechter et al., 1958; Turner et al., 2012; Jun and Taheri-Araghi, 2015). However, some modes of cellular growth rate regulation in bacteria escape this general rule (Basan et al., 2015; Si et al., 2017). Early work in eukaryotic microbes such as fission yeast studied the steady-state relationship between average protein mass per cell and population growth rate for different conditions, and concluded that “nutritional reduction of growth rate leads to a decrease in cell size”(Fantes and Nurse, 1977) in line with the generic rule. Fission yeast is an attractive eukaryotic model organism to investigate the relationship between cell size, cell geometry and growth conditions. Specifically, it is rod-shaped, it has a small genome, it is genetically easy to manipulate, and its cell cycle is very well studied (Hoffman et al., 2015). Molecular studies have identified pathways connecting nitrogen sensing to the cell-cycle machinery controlling the G2/M transition, resulting in rapid acceleration or inhibition of mitosis commitment upon sudden changes of the nitrogen source in the medium (Carlson et al., 1999; Petersen and Nurse, 2007; Chica et al., 2016). However, it has not yet been reported how different features of the cell geometry, such as volume, length, width or surface-to-volume ratio, are regulated across different types of growth rate modulation (Gu and Oliferenko, 2021; Kellogg and Levin, 2022).

To revisit this problem, we designed an experimental setup combining continuous culturing with single-cell microscopy to modulate and quantify the steady-state growth rate and the corresponding cell geometry at division (Figure panel A, Methods). Media inflow was controlled to maintain the optical density of the culture near a setpoint value. Turbidostat cultures are advantageous over chemostat cultures because cell density can be kept independent from the composition of the growth medium. This prevents confounding effects on cell physiology caused by accumulation of waste metabolites in the growth medium at high cell densities. Furthermore, controlling cellular growth in turbidostats also avoids the need to externally adapt the flow rate to each growth medium to prevent washout; rather, the propagation rate is entirely set in response to processes internal to the cells under investigation. As in other continuous culturing setups, turbidostat cultures reach steady state after growing in identical conditions for >10 generations.

To identify mitotic cells, we used cells with the septum protein Mid2 tagged with GFP (Tasto et al., 2003). To accurately quantify the geometry of those cells, we acquired multiple z-stacks and used a semi-automated image analysis pipeline to extract individual cell contours. These contours were then divided into small segments perpendicular to their long axis, enabling us to compute cell length, width, surface area, and volume, assuming rotational symmetry (Figure panel B, Methods). This approach remains accurate even when cell width varies along the length axis, such that the cell shape deviates from a perfect spherocylinder.

### Change of cell length, surface area and volume at division with growth rate under various growth modulations

We obtained 17 turbidostat cultures covering three types of growth rate modulations: low nitrogen (10– 50 mg/L of ammonium chloride (NH_4_Cl) in what was otherwise standard EMM2 minimal medium), low glucose (0.5-2.0 g/L of glucose in otherwise EMM2), and protein synthesis inhibition (by adding growth-inhibitory doses of 0.5–3.0 mg/L cycloheximide in EMM2 medium). Measurements of cell geometry are compared to the population growth rate (Figure). In addition, we include data obtained from cultures where different sources of nitrogen were used (for reference, NH_4_Cl at 5 g/ml or 93.5 mM, which is equal to EMM2 medium; alternatively proline, serine, phenylalanine, glycine, and tryptophan all at 20 mM). For these six cultures, glucose and the respective nitrogen source were available in abundance and protein synthesis was not inhibited. Of note, growth rates as well as proteomes and transcriptomes data of cells from these cultures were recently published elsewhere (Kleijn et al., 2022). Figure panels show the average values of the volume (Panel D), surface-to-volume ratio (Panel E), surface area (Panel F), cell length (Panel G), and cell width (Panel H) plotted against the growth rate. Panel C shows an expected relationship between surface-to-volume ratio and cell width, highlighting consistency of our image analysis and cell geometry quantification across all growth conditions.

We first looked at glucose limitation. There, the three size metrics (cell length, surface area and volume) decreased as growth rate decreased, in agreement with the nutritional control model of cell size regulation (Fantes and Nurse, 1977; Petersen and Nurse, 2007). Next, we assessed growth in conditions of ammonium limitation. Here the picture was more complex. Under moderate ammonium limitations (NH_4_Cl concentrations between 25 and 50 mg/L), cell length was increased compared to unperturbed cultures. However, in this regime the surface area was not significantly different compared to the base medium. In contrast, when looking at the whole range of ammonium concentrations between 10 and 50 mg/L, the three size metrics all varied coordinately and approximately linearly with the growth rate.

Strikingly, while the relations between the growth rate and cell length and width were different between glucose and ammonium limitations, the dependence of the cell volume on growth rate appeared identical across the entire range of concentrations. Because cell geometry differed clearly between the two modulations (cells were longer and thinner under low ammonium compared to low glucose), this suggests the existence of a geometry-independent link between cell size at division and growth rate at low nutrient concentrations.

Next, we analysed fission yeast growth on a series of alternative nitrogen sources at saturating concentration (Carlson et al., 1999). The growth rates and size values observed in this study were consistent with previously reported growth rate (Carlson et al., 1999) and protein content measurements (Fantes and Nurse, 1977; Petersen and Nurse, 2007). Specifically, the protein content of cells grown on proline was considerably smaller than that of cells grown on serine or ammonium chloride in chemostats (Fantes and Nurse, 1977). However, in contrast to growth on limited ammonium or glucose, we did not observe any clear trend between the cell size metrics and the growth rate. Notably, proline was the only nitrogen source whose cell volumes were on the regression line defined by the low-ammonium and low-glucose conditions; for the other four sources the cell volume was ∼2-3x larger than in low-ammonium at equivalent growth rates. These results indicate that the nutritional control of size is qualitatively different depending on whether we consider nutrient quality or nutrient abundance. At steady-state, the ‘smaller in poorer medium’ rule only applies to nutrient abundance and not nutrient quality.

We next examined cell geometry variations under growth rate modulations caused by inhibiting translation with cycloheximide. We observed only slight changes of cell size metrics as the growth rate decreased (small decreases in length and small increases in surface area and in volume). These results are consistent with the effect of translation inhibition on steady-state cell size (using chloramphenicol) in the bacterium *E. coli* (Basan et al., 2015; Si et al., 2017). We note that our data covers a narrower growth rate range for cycloheximide modulation than the others, because larger cycloheximide concentrations induced significant morphological aberrations, including multi-septated cells.

Finally, we looked at regulation of the cell’s surface-to-volume ratio. For rod-shaped fission yeast cells, the surface-to-volume ratio remains mostly constant during the cell cycle and is dictated predominantly by cell width (Mitchison and Nurse, 1985; Odermatt et al., 2021). Any control mechanism regulating the surface-to-volume ratio could therefore act independently from the cell volume at division. We computed the average surface-to-volume ratio of mitotic cells for all growth rate modulations (Figure panel E). We observed either negative (glucose and ammonium titration), positive (translation inhibition) or no (nitrogen source quality) correlation of the surface-to-volume ratio with the growth rate. In low ammonium concentrations, cells showed markedly increased surface-to-volume ratio. This increase in surface area relative to cell volume might favor the uptake of ammonium by membrane transporters located on the cell surface. Contrasting with this, in low glucose concentrations, cells only moderately increased their surface-to-volume ratio. Under translation inhibition, the surface-to-volume ratio moderately decreased with the growth rate. In summary, the surface-to volume ratio and cell volume relate to the growth rate in different ways.

## Discussion

Established literature of pombe size regulation by the environment identified nutritional control via the TOR pathway as a main regulatory mechanism (Petersen and Nurse, 2007; Yanagida et al., 2011; Gonzalez and Rallis, 2017). In this context, we report an account of the changes in cell geometry that occur during growth on media limited in high-quality nutrient, during growth on low quality nutrients, or in response to a translation inhibitor. For this we used turbidostat cultures and geometry-aware cell size quantification with single-cell microscopy. Overall, we find that the relation between cell geometry and the growth rate tends to vary depending on the mode of growth rate modulation.

This work also uncovers a linear size law (Figure D): under low nutrient conditions in steady-state, the cell volume and growth rate are linearly correlated regardless of whether growth is limited by carbon or nitrogen availability. We therefore propose that while surface-to-volume ratio is regulated specifically in response to the type of limiting nutrient (Figure E), the mechanisms controlling division are coupling cell volume (and not surface area or cell length) to growth rate. We speculate that such coupling might serve the evolutionary purpose of preparing cells to total growth arrest, a state where it is advantageous to reach a small size per unit of genome to reduce cellular maintenance costs. On the contrary, the lack of correlation between cell size and growth rate when growth rate is modulated by the quality (and not abundance) of the nitrogen source can be rationalized by the fact that because current nutrients are abundant, it is unlikely that a total growth arrest will occur soon.

It has been reported that fission yeast size homeostasis is controlled by the Cdr2 nodal density, which indirectly measures the cell surface area (Pan et al., 2014; Facchetti et al., 2019). In these experiments, cell mutants with different widths were studied in reference media, and a cryptic volume control was found. Other recent studies have proposed Cdc25 and/or Cdc13 as reporters of cell volume (Keifenheim et al., 2017; Curran et al., 2022; Miller et al., 2022; Bashir et al., 2023). We speculate that the observed volume scaling with nutrient titration (panel D) may be regulated by Cdc25 and Cdc13 rather than Cdr2 that senses cell surface.

We also observe that the surface-to-volume ratio is increased with nutrient limitation (panel E), facilitating nutrient import. However, under carbon limitation it is less strongly modulated than under nitrogen limitation. A plausible explanation for this comes from the observation that the cell surface is mainly composed of carbon, and carbon limitation could thereby limit the increase of the surface-to-volume ratio. Mechanistic models of cell size control and cellular resource allocations could in principle be used to explain this behavior, as we have recently done for bacteria (Bertaux et al., 2020; Kleijn et al., 2023).

## Methods

### Strains and media

Wild type fission yeast with GFP-tagged Mid2 was obtained from the Gould laboratory (Tasto et al., 2003).

All growth media were derived from Edinburgh minimal media (EMM2, (Petersen and Russell, 2016)). Four types of modifications were made to the standard EMM2 medium:

1. Decreased concentrations of ammonium chloride (chosen as 10, 15, 20, 25, 30, 40, and 50 mg/L; the concentration in EMM2 is 5.0 g/L or 93.5 mM).
2. Decreased concentrations of glucose (set to 0.50, 0.75, 1.0, 1.5, and 2.0 g/L; in EMM2 this is 20 g/L or 111 mM).
3. Added the translational inhibitor cycloheximide in nonlethal quantities (concentrations of 0.50, 1.0, 2.0, and 3.0 mg/L, not present in standard EMM2).
4. Saturating concentrations (20 mM) of five alternative amino acids (proline (Pro), serine (Ser), phenylalanine (Phe), glycine (Gly), and tryptophan (Trp)) were used instead of the standard ammonium chloride (Amm).

### Cell culture and imaging

The turbidostat culturing procedures have been previously described and characterized in detail (Kleijn et al., 2022). Briefly, cells were first precultured on YES agar plates, then twice in flask cultures at 32°C to around 5×10^6^ cells/ml, before inoculating the turbidostat cultures at 0.5–1.0×10^6^ cells/ml. These cultures were placed in syphoning turbidostat devices at 32°C (Takahashi et al., 2015). This setup enabled us to control media flow every 30s to maintain OD_600_ at a fixed set point, corresponding to 3–5×10^6^ cells/ml depending on the growth conditions. To ensure that steady state was reached, the cultures were left in the turbidostats for between 30 and 80 hours depending on the growth medium, corresponding to at least 10 cell generations.

For the cultures limited by the quality of abundant nitrogen source, the turbidostat set point was maintained at OD_600_=0.40. Furthermore, culture growth rates were measured approximately halfway through each experiment, by diluting the cultures twofold and fitting exponential growth to the regrowth phase back to OD_600_=0.40 (Kleijn et al., 2022). For the titration experiments, the set point was at OD_600_=0.30 and growth rates were measured at the end of the culturing directly following cell harvest, by releasing the threshold and fitting an exponential to the subsequent increase in cell density.

Cell samples were collected, loaded on imaging slides (ibidi, μ-Slide VI 0.4), and transferred to the microscope to be imaged. Slides were imaged on an Olympus IX-70 for brightfield and Mid2-GFP fluorescence. Exposure times were 30ms for brightfield images and 1000ms for GFP images. In both the brightfield and GFP channels, six images were obtained with focal depth 1.5 μm apart.

### Image analysis

Images were analysed with custom MATLAB scripts using the Image Processing Toolbox. Cells undergoing septation were manually selected based on the intracellular location of the Mid2-GFP stain. For septated cells, a two-dimensional cross-sectional contour was extracted from the brightfield image. For cells that passed a visual inspection of their cross-sections, cell shape data was inferred as follows (Figure panel B). First, the length *L* of the cell was taken to be the distance between the two cell extremities. Then, the contour was discretized into *N*=50 segments perpendicular to the length axis of the cell, and for each segment *i* the width *w*_*i*_ was calculated. The cell width *w* was taken as the average width of the slices:

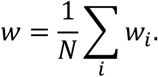

Assuming cylindrical symmetry for each individual segment, the total volume *V* and surface *S* of the cell were then approximated as

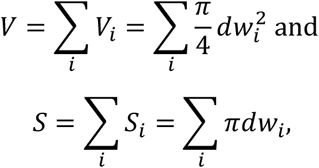

with 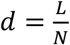 the height of each of the cylinders.

## Code and data availability

A supplementary table with single cell data is provided with the paper, and R code generating the figure will be shared on GitHub upon publication.

